# Long-term recoveries of forest ecosystem components after shallow landslides were asynchronized among slope positions

**DOI:** 10.1101/2022.11.07.515411

**Authors:** Wataru Hotta, Junko Morimoto, Seiji Yanai, Yoshitaka Uchida, Futoshi Nakamura

**Author notes:** **Mailing address**, Wataru Hotta, Junko Morimoto, Seiji Yanai, Yoshitaka Uchida, Futoshi Nakamura. **Author contributions**, WH and JM conceptualized this study. SY provided information on past landslides. WH performed the field survey, data processing, and data analyses. WH and YU performed, soil analyses. WH wrote the original draft. All authors contributed critically to the drafts and gave final approval for publication. **Corresponding author**: W. Hotta, Postal Address: Graduate School of Agriculture, Hokkaido University, Kita 9, Nishi 9, Kita ku, Sapporo, Hokkaido 060-8589, Japan.

## Abstract

Landslides are a common disturbance in mountainous areas of the world. Transporting and accumulating landslide debris, i.e., disturbance legacies, such as coarse woody debris (CWD), vegetation patches, and surface soils, generate a heterogeneous environment along slopes (zones), which are suggested to affect forest recovery. However, the long-term changes in forest ecosystems after landslides remain unknown, particularly zone-dependent change patterns. We aimed to reveal the differences in the changes in live trees, understory vegetation, CWD, and soils among zones by surveying forests with landslide ages (years since the landslide) ranging from 3 to 74 years and reference stands. The increase in live tree aboveground biomass occurred at a faster rate at the lower part of the slopes where the disturbance legacies were rich and surface soils were stabilized due to the smaller slope angle. Chronological patterns of understory vegetation amounts were determined by the differences in disturbance legacy richness and the timing of subsequent canopy closure. The amounts and decay-class diversity of CWD were determined by the differences in legacy richness and mortality through stand development. These zone-dependent chronological changes influenced litter production and determined the recovery rates of surface soil carbon stocks and nitrogen contents. The increase in the dominance rates of forest herbaceous species was faster in the lower part of the slopes due to the faster surface soil development and canopy closure. Our results illustrate that long-term forest ecosystem succession and recovery after landslides occurs more rapidly at the lower parts than at the upper parts of slopes.

**Highlights:**

- Rich landslide legacies and low slope angles promoted vegetation and soil recovery
- Legacy richness and timing of canopy closure determined changes in the understory
- Landslide mortality and stand development mortality determined changes in dead wood

## INTRODUCTION

Natural disturbances are one of the major factors affecting the fundamental processes in forest ecosystems, such as plant growth, mortality, and decomposition of litter and soils (Turner 2010). Recoveries of multiple ecosystem components need to be evaluated to understand forest ecosystem recovery after disturbances because forest ecosystems are the associative systems of live trees, coarse woody debris (CWD), understory vegetation, and soils. Live trees account for most forest biomass and are commonly used as indicators of ecosystem recovery after disturbances (Roberts 2004). Organic matter assimilated in live trees transfers CWD or surface soil pools through mortality and falling leaves. CWD in its diverse decay stages plays important roles as habitats for insects or animals and regeneration sites for trees (Vanderwel and others 2006; Weaver and others 2009) and is eventually transferred to soils (Harmon and others 1986). Understory vegetation accounts for less than 1% of forest biomass; however, it accounts for over 90% of plant species in forests (Gilliam 2007). Herbaceous species, which account for most of the understory vegetation, contain higher concentrations of nutrients such as nitrogen and are decomposed more rapidly than woody species (Muller 2003). Soils are formed by the accumulation and decomposition of litters derived from live trees and understory vegetation, which play fundamental roles in the provision of vital nutrients for plant growth. However, once forests are disturbed, several processes in forest ecosystems are altered, e.g., massive live tree die-off, changes in plant species composition due to improvement of light conditions, increasing decomposition rates of CWD and soils, etc. (Zhang and Liang 1995). These influences vary widely among disturbance types and intensities (Turner 2010). Thus, it is important to understand how each forest ecosystem component links and recovers or changes for a comprehensive understanding of forest ecosystem recovery after disturbances.

Landslides are one of the major natural disturbances in mountainous areas of the world (Restrepo and others 2009; Veblen and others 1992) and are predicted to occur more frequently and intensely in the future because of the increase in heavy rain events, which is the major trigger of shallow landslides due to climate change (Lateltin and others 1997). Long-term recoveries of live trees, understory vegetation, and soils after landslides have been studied. It has been reported that it takes more than 25-120 years for stand basal area recovery (Pandey and Singh 1985; Reddy and Singh 1993; Restrepo and others 2003); on the other hand, soil carbon stocks and nitrogen content recover within 35-120 years (Pandey and Singh 1985; Reddy and Singh 1993). The amount of understory vegetation has been reported to gradually increase with stand ages up to 120 years after landslides in sync with relatively slower tree biomass recovery (Reddy and Singh 1993), although it gradually decreased with stand ages up to 427 years due to canopy closures, according to a study focused on recovery after harvesting (Jules and others 2008). In contrast, long-term changes in CWD after landslides are poorly understood. Studies focusing on disturbances other than landslides reported that changes in the amount of CWD with stand age showed “U-shaped temporal patterns” after various disturbance types (Brassard and Chen 2008; Krankina and Harmon 1995; Spies and others 1988); i.e., the amount of CWD was greater in the young and old stages but relatively less in the intermediate stage in forest developments. These patterns are shaped by the combination of carryover, disturbance mortality, and input from new stands (Spies and others 1988). However, this information cannot be simply applied to forest ecosystem dynamics after landslides dependent on locations along the slope because landslides shape relatively distinct zones by transporting and accumulating undisturbed vegetation patches, CWD, and soils (Allen and others 2020; Restrepo and others 2003).

### No study has evaluated the differences in the long-term (over 50 years) changes in multiple forest ecosystem components among the zones generated by landslides

Each zone generated by a landslide has completely different characteristics based on the amount of disturbance legacy (i.e., undisturbed vegetation patches, soils and CWD transported from the upper to lower part of slopes; Johnstone and others 2016) and the physical environment such as slope angle and soil moisture content: the initiation zone is the starting point of landslides and is characterized by a steep slope and infertile soils; the deposition zone is the end part of landslides and legacies, which existed on slopes, are accumulated here; and the transport zone is the transition zone between the two zones mentioned above (Martin and others 2002, Walker and Shields 2012). Because of this heterogeneity, previous studies suggested that tree, herb, and soil recoveries would be slower on the upper parts of slopes because of lower soil stabilities and fertilities due to higher slope angles (Buma and Pawlik 2021; Guariguata 1990; Wicke and others 2003). However, existing studies considering zones have mainly focused on a single ecosystem component or on short-term (less than 10 years) responses of ecosystems, although several hundred years are needed for forest development, and ecosystem recovery should be evaluated by investigating the dynamics of multiple ecosystem components, which resulted that we still have a limited knowledge of the ecosystem recovery after landslides.

This study aimed to reveal how the long-term (over 70 years) change patterns of multiple forest ecosystem components after shallow landslides differed among the zones generated by landslides. We hypothesized that the recovery rates of live tree aboveground biomass (AGB), surface soil carbon stocks and nitrogen content (CN) and succession of species composition occurred at a faster rate in the deposition zone than the transport and the initiation zones due to the richer disturbance legacies and the soil stability at the lower slope angle positions. Additionally, the chronological changes in understory vegetation and CWD were hypothesized to be different among the zones dependent on the amounts of disturbance legacies remaining and the subsequent stand development.

## MATERIALS & METHODS

### Study area

The study area was located in mixed forests in the transition zone of the cool-temperate and boreal areas in southern Hokkaido, northern Japan (Figure 1). The annual mean temperature and precipitation are 7.0 °C and 1,028.4 mm, respectively, at the meteorological observation station in Atsuma (42°43’48”N, 141°53’18”E; at 20 m a.s.l.), and 7.6 °C and 983.7 mm, respectively, at the meteorological observation station in Hidaka-Monbetsu (42°29’48”N, 142°3’12”E; at 47 m a.s.l.) (Japanese Meteorological Agency 2022; average of 1991-2020). The altitude above sea level is 40–200 m. Because of the low temperature (below 0 □) and relatively thin snow cover (21-69 cm at maximum) during winter, freeze-thaw action causes soil instability. The dominant parent material consists of Tertiary mudstone and conglomerate, Quaternary terrace deposits, volcanic ash, and pumice (Geological Survey of Japan, AIST 2022). The dominant soil types are Andosols and partly volcanogenous Regosol (IUSS Working Group WRB 2015). The forest canopy around landslides is dominated by cool-temperate broadleaves such as *Tilia japonica* (Miq.) Simonk, *Acer pictum* Thunb., and *Quercus crispula* Blume var. *crispula*.

**Figure 1.**
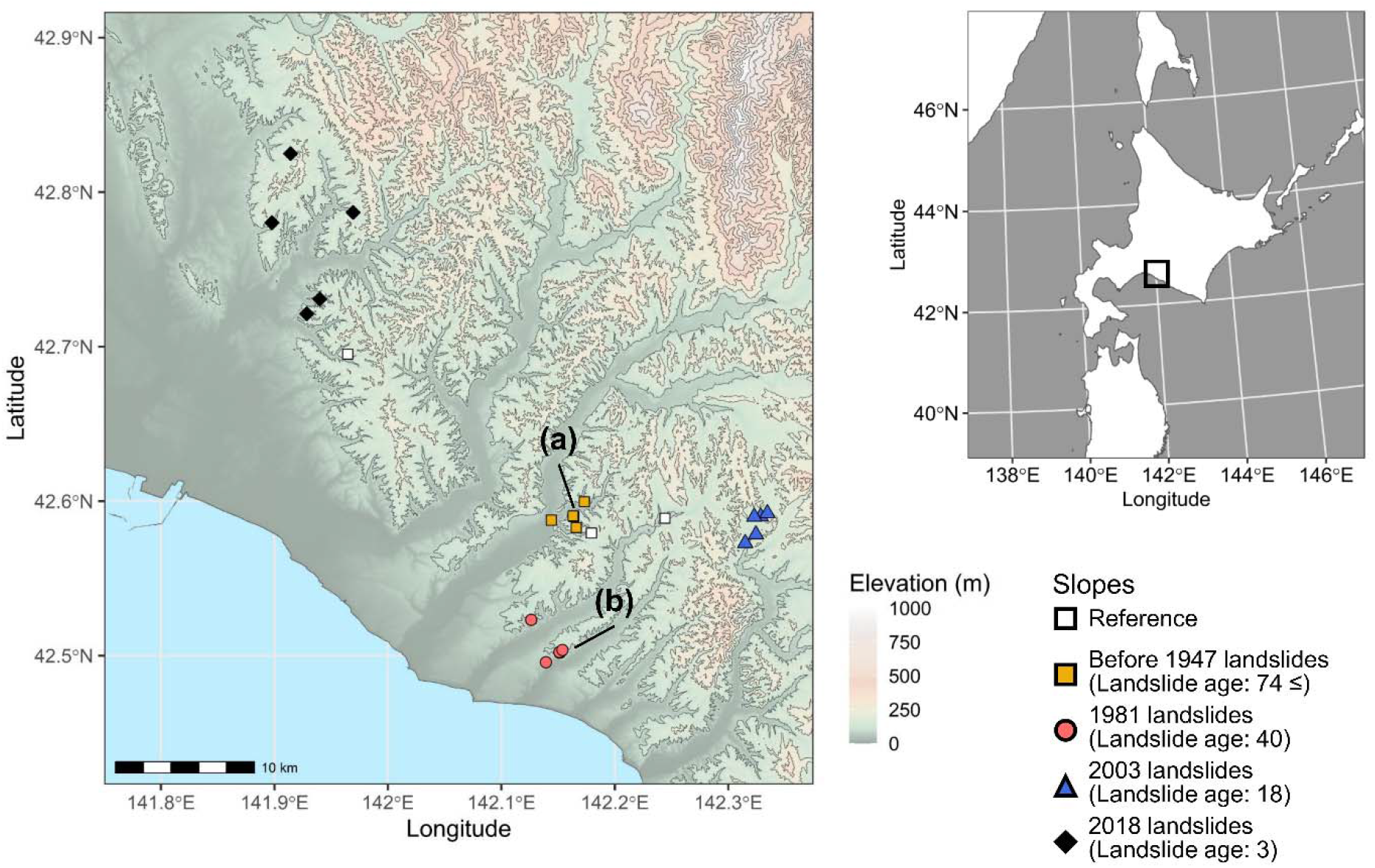
Map of the study area. Right: Map of northern Japan and its surroundings. The black rectangle indicates the range of the detailed map on the left. Left: The detailed map of the study area. The points indicate the location of the study landslides and slopes. (a) Two slopes where landslides occurred before 1947. (b) Three slopes where landslides occurred in 1981.

### Placement of plots by reviewing the landslide disturbance history

We used chronosequence approaches to estimate the long-term forest ecosystem changes. We chose five landslides for each landslide age (years since the landslide; 3, 18, 40, and over 74 years post-landslides; i.e., landslides occurred in 2018, 2003, 1981, and before 1947) and three reference stands that are considered not to be disturbed over 100 years. The areas of landslides ranged from 0.07 to 0.29 ha. We selected suitable landslides and stands (Figure 1) by reviewing the time series of aerial photographs and the records of previous studies (Yanai and Igarashi 1990; Sato and others 2013). We divided each landslide or stand into three zones: initiation, transport, and deposition for landslides (Restrepo and others 2009; Walker and others 2009) and upper, middle, and lower for reference stands (Appendix 1). In each zone, we established one plot of 15 m × 15 m for surveying living trees with a height larger than 1.3 m and CWD, four quadrats of 2 m × 2 m for surveying environmental conditions and the understory vegetation, which included all nontree vascular plants and trees with a height smaller than 1.3 m, and one quadrat of 0.25 m × 0.25 m for sampling organic (O) layer and mineral soils. We did not establish plots and quadrats in the deposition zone of landslides over 74 years ago because roads and farmland were newly built in their deposition zones, and we could not find suitable survey areas.

### Field data collection

For the living trees with a height larger than 1.3 m, we recorded species and measured the diameter at breast height (DBH) with a tree caliper (when DBH was larger than 46 cm, it was measured with a measuring tape) and tree heights with a TruPulse 360 rangefinder (Laser Technology Inc., USA).

We classified CWD into three types: (1) snags: standing dead trees taller than 1.5 m; (2) downed logs: dead trees that lay on the ground and did not have roots and that lay on the ground with roots and stems longer than or equal to 1.5 m; and (3) stumps: standing dead trees shorter than 1.5 m and dead trees laying on the ground with roots and stems shorter than 1.5 m (Forestry and Forest Products Research Institute 2016). Snags with DBHs larger than 5 cm were targeted for recording whether conifers or broadleaves and the decay class and measuring DBH and heights. Downed logs with the diameter of either end larger than 10 cm were targeted for recording whether conifers or broadleaves and the decay class and measuring diameters of both ends and lengths. We targeted stumps with a diameter of the upper end larger than 5 cm; however, no such stumps were observed in any of the plots. We used the five-level decay classification system (i.e., 1, 2, 3, 4, and 5, where 1 is the newest and 5 is the oldest) described in Graham and Cromack (1982). The decay class of CWD was identified based on the morphological characteristics and the depths of the knife sticks (Fukasawa 2012; Heilmann-Clausen 2001).

For the understory vegetation, we identified species and measured the coverage and the maximum height of each species. Environmental variables such as soil moisture content, slope angle, and O layer depth were measured. Time domain reflectometry (HydroSense, Campbell Scientific, Inc.) was used to measure the water content of the soil by volume. The soil water content was measured on a day following three continuous sunny days in mid-October 2021 and measured three times at 5 cm beneath the surface in each quadrat. The slope angle and O layer depth were measured at the center of each quadrat.

We sampled the O layer and mineral soils in 0.25 m × 0.25 m quadrats. We classified the O layer into the Oi layer, the Oa/Oe layer (the Oi layer contains fresh leaf litter, but the Oa/Oe layer contains well-decomposed litter), and twigs and then sampled all of the O layer within the quadrats. We sampled the surface mineral soils (0-5 cm depth) in each quadrat by using cylindrical cores (100 mL) and measured the bulk density. We also sampled approximately 200 g of surface mineral soils (0-5 cm depth) by shovels to analyze the carbon and nitrogen concentrations.

All surveys were conducted from July to October 2021.

### Calculations of live tree AGB, CWD mass, and the biomass index of understory vegetation

Live tree AGB was calculated using species-specific equations. Downed log mass was calculated using the conic-paraboloid formula (Fraver and others 2007) and the decay class specific volume density. Snag mass was calculated by approximating their shape to cones. Please see Appendix 2 for details on the mass calculations. The biomass index of the understory vegetation was calculated as the sum of the coverage of each species multiplied by the maximum height of each species (Olson and Martin 1981; Suchar and Crookston 2010).

### Soil analysis

O layer samples were dried in an oven (70 °C, 48 h), and their dry weights were measured. Then, we ground them by using a Wiley mill (W-100S, IRIE SHOKAI Co., Ltd., Japan) and a ball mill (Sample Mill TI-100, CMT Co., Ltd., Japan). Finally, the carbon and nitrogen concentrations of each sample were measured using a CHNS/O analyzer (PE-2400, PerkinElmer, USA).

Mineral soil samples for measuring bulk densities were dried, and we classified each mineral soil sample for measuring bulk density into fine soil particles, gravel, and roots by passing them through a 2 mm sieve. After drying, the dry weight of the fine soil was measured. The bulk density was calculated according to the dry weight of fine soils divided by the volume (i.e., 100 mL).

The mineral soils collected for measuring carbon and nitrogen concentrations were air dried and then passed through a 2 mm sieve followed by a 500 μm sieve. The carbon and nitrogen concentrations of each sample were measured using a CHNS/O analyzer (PE-2400, PerkinElmer, USA).

### Statistical analysis

Differences in environmental variables, live tree AGB, CWD mass, the biomass index of understory vegetation, surface soil carbon stock and nitrogen content, including the O layer and mineral soils among landslide ages and zones, were tested with a generalized linear mixed model (GLMM; *lmerTest* package) and multiple comparisons (*multcomp* package; *p* adjustment methods: Benjamini–Hochberg method (Benjamini and Hochberg 1995)). The explanatory variable was the combination of landslide ages and zones. The random effect was the landslide identity, and the error structure was a normal distribution.

Differences in the species composition of living trees and understory vegetation were tested with permutational analysis of variance (PERMANOVA; *vegan* package; Oksanen and others 2020) and multiple comparisons. The explanatory variable was the combination of landslide ages and zones. The response variables were the basal area of each living tree species and the vegetation cover of each plant species. Additionally, the non-metric multidimensional scaling (NMDS) was conducted to visualize differences in species compositions among the combinations of landslide ages and zones and to identify environmental factors determining species compositions (*vegan* package; Oksanen and others 2020). We considered slope angle, soil moisture content, surface soil nitrogen content, and O layer depth as environmental factors in NMDS. For the understory analysis, AGB was additionally considered as an environmental factor. If the *p* value was lower than 0.05 in the analysis using the function *envfit*, then we regarded the relevant factor as the determining factor of species compositions. Note that the plots where zero or only one individual was observed were excluded from analyses of live tree species composition; i.e., the plots in landslide ages 3 and 18 were excluded.

We did not treat landslide ages and zones as independent explanatory variables because of the incomplete design due to the lack of data of the deposition zone in landslides over 74 years. All statistical analyses were conducted in R 4.0.2 (R core team 2020). Additionally, we classified live tree species by shade and desiccation tolerance. Herbaceous species were classified into three species types, i.e., open land species, generalist species, and forest species, and three moisture types, i.e., dry, moderate, and wet.

## RESULTS

### Environmental characteristics of landslide ages and zones

The differences in slope angles were mainly explained by zones because slope angles were significantly different among the zones for almost all landslide ages (Table 1). The differences in O layer depths were mainly explained by landslide age; however, significant differences among the zones were observed for landslide age 18 (Table 1). The differences in soil moisture contents were mainly explained by zones, although there were some minor variations by landslide age (Table 1).

**Table 1.**
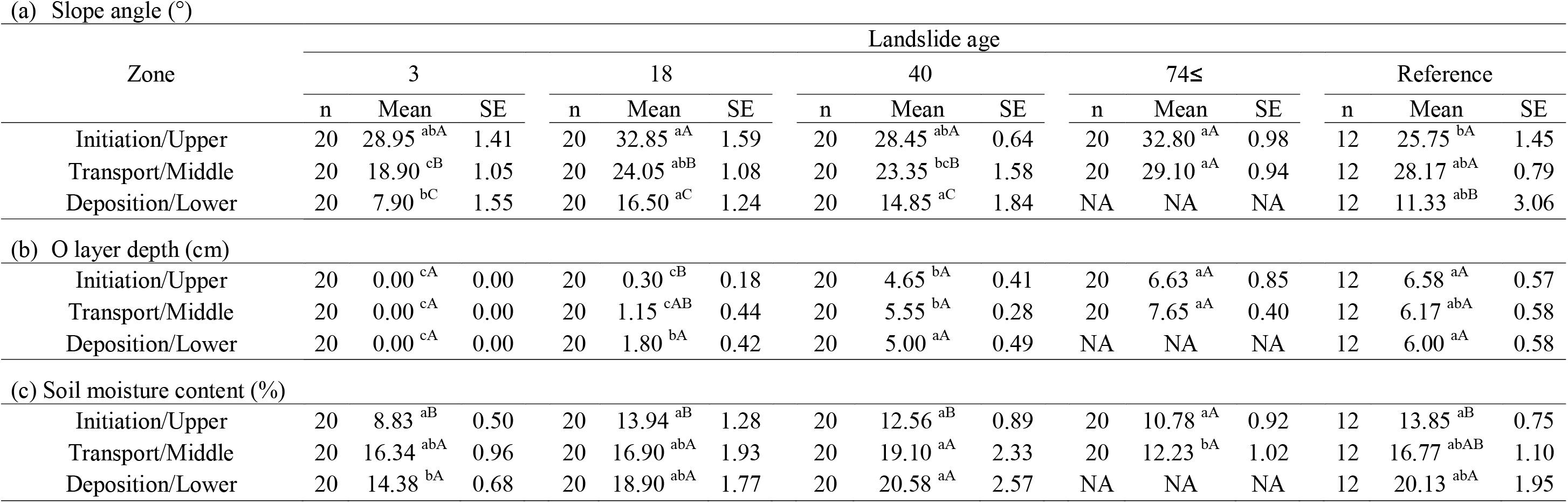
Environmental variables by landslide ages and zones. Different lowercase letters next to the values indicate significant differences between landslide ages within each zone (p < 0.05) according to multiple comparisons. Different capital letters next to the values indicate significant differences between zones within each landslide age (p < 0.05) according to multiple comparisons.

| Zone | Landslide age |  |  |  |  |  |  |  |  |  |  |  |  |  |  |
| --- | --- | --- | --- | --- | --- | --- | --- | --- | --- | --- | --- | --- | --- | --- | --- |
|  | 3 |  |  | 18 |  |  | 40 |  |  | 74≤ |  |  | Reference |  |  |
|  | n | Mean | SE | n | Mean | SE | n | Mean | SE | n | Mean | SE | n | Mean | SE |
| Initiation/Upper | 20 | 28.95<sup>abA</sup> | 1.41 | 20 | 32.85<sup>aA</sup> | 1.59 | 20 | 28.45<sup>abA</sup> | 0.64 | 20 | 32.80<sup>aA</sup> | 0.98 | 12 | 25.75<sup>bA</sup> | 1.45 |
| Transport/Middle | 20 | 18.90<sup>cB</sup> | 1.05 | 20 | 24.05<sup>abB</sup> | 1.08 | 20 | 23.35<sup>bcB</sup> | 1.58 | 20 | 29.10<sup>aA</sup> | 0.94 | 12 | 28.17<sup>abA</sup> | 0.79 |
| Deposition/Lower | 20 | 7.90<sup>bC</sup> | 1.55 | 20 | 16.50<sup>aC</sup> | 1.24 | 20 | 14.85<sup>aC</sup> | 1.84 | NA | NA | NA | 12 | 11.33<sup>abB</sup> | 3.06 |
| (b) O layer depth (cm) |  |  |  |  |  |  |  |  |  |  |  |  |  |  |  |
| Initiation/Upper | 20 | 0.00<sup>cA</sup> | 0.00 | 20 | 0.30<sup>cB</sup> | 0.18 | 20 | 4.65<sup>bA</sup> | 0.41 | 20 | 6.63<sup>aA</sup> | 0.85 | 12 | 6.58<sup>aA</sup> | 0.57 |
| Transport/Middle | 20 | 0.00<sup>cA</sup> | 0.00 | 20 | 1.15<sup>cAB</sup> | 0.44 | 20 | 5.55<sup>bA</sup> | 0.28 | 20 | 7.65<sup>aA</sup> | 0.40 | 12 | 6.17<sup>abA</sup> | 0.58 |
| Deposition/Lower | 20 | 0.00<sup>cA</sup> | 0.00 | 20 | 1.80<sup>bA</sup> | 0.42 | 20 | 5.00<sup>aA</sup> | 0.49 | NA | NA | NA | 12 | 6.00<sup>aA</sup> | 0.58 |

**Table 1 Environmental variables by landslide ages and zones.**
|  |  |  |  |  |  |  |  |  |  |  |  |  |  |  |  |
| --- | --- | --- | --- | --- | --- | --- | --- | --- | --- | --- | --- | --- | --- | --- | --- |
| Initiation/Upper | 20 | 8.83<sup>aB</sup> | 0.50 | 20 | 13.94<sup>aB</sup> | 1.28 | 20 | 12.56<sup>aB</sup> | 0.89 | 20 | 10.78<sup>aA</sup> | 0.92 | 12 | 13.85<sup>aB</sup> | 0.75 |
| Transport/Middle | 20 | 16.34<sup>abA</sup> | 0.96 | 20 | 16.90<sup>abA</sup> | 1.93 | 20 | 19.10<sup>aA</sup> | 2.33 | 20 | 12.23<sup>bA</sup> | 1.02 | 12 | 16.77<sup>abAB</sup> | 1.10 |
| Deposition/Lower | 20 | 14.38<sup>bA</sup> | 0.68 | 20 | 18.90<sup>abA</sup> | 1.77 | 20 | 20.58<sup>aA</sup> | 2.57 | NA | NA | NA | 12 | 20.13<sup>abA</sup> | 1.95 |

### Initial ecosystem conditions after the landslide

The amounts of understory vegetation and CWD were significantly larger in the deposition zone than in the initiation and transport zones with landslide age 3, which is the earliest age after a landslide in this study (Table 2). Live tree AGB and surface soil CN were not significantly different among the zones.

**Table 2.** Amounts of each forest ecosystem component for landslide age 3. Different capital letters next to the values indicate significant differences in the amount of each forest ecosystem component among the zones (p < 0.05) according to multiple comparisons. AGB: aboveground biomass; CWD: coarse woody debris.

| Zone | Live trees AGB<br>(Mg ha<sup>-1</sup>) |  |  | Understory<br>vegetation<br>biomass index<br>(m<sup>-2</sup>) |  |  | CWD<br>(Mg ha<sup>-1</sup>) |  |  | Surface soil<br>carbon stock<br>(MgC ha<sup>-1</sup>) |  |  | Surface soil<br>nitrogen content<br>(MgN ha<sup>-1</sup>) |  |  |
| --- | --- | --- | --- | --- | --- | --- | --- | --- | --- | --- | --- | --- | --- | --- | --- |
|  | n | Mean | SE | n | Mean | SE | n | Mean | SE | n | Mean | SE | n | Mean | SE |
| Initiation | 5 | 0.00<sup>A</sup> | 0.00 | 20 | 1.35<sup>B</sup> | 0.32 | 5 | 0.00<sup>B</sup> | 0.00 | 5 | 5.25<sup>A</sup> | 1.53 | 5 | 0.53<sup>A</sup> | 0.13 |
| Transport | 5 | 0.00<sup>A</sup> | 0.00 | 20 | 3.61<sup>B</sup> | 0.47 | 5 | 3.71<sup>B</sup> | 2.54 | 5 | 3.31<sup>A</sup> | 0.76 | 5 | 0.71<sup>A</sup> | 0.08 |
| Deposition | 5 | 0.00<sup>A</sup> | 0.00 | 20 | 19.86<sup>A</sup> | 2.81 | 5 | 14.12<sup>A</sup> | 4.54 | 5 | 4.37<sup>A</sup> | 1.06 | 5 | 0.47<sup>A</sup> | 0.08 |

### Live trees

Live tree AGB increased regardless of zone; however, the increase in AGB was slower in the initiation zone than in the other zones (Figure 2a). AGB 18 and 40 years after landslides was significantly different in the transport and deposition zones (p < 0.05) but not in the initiation zone. AGB over 74 years after landslides was slightly significantly different between the initiation and transport zones (p < 0.10). Additionally, AGB over 74 years after landslides was still significantly different in all zones from that in the reference stands.

**Figure 2.**
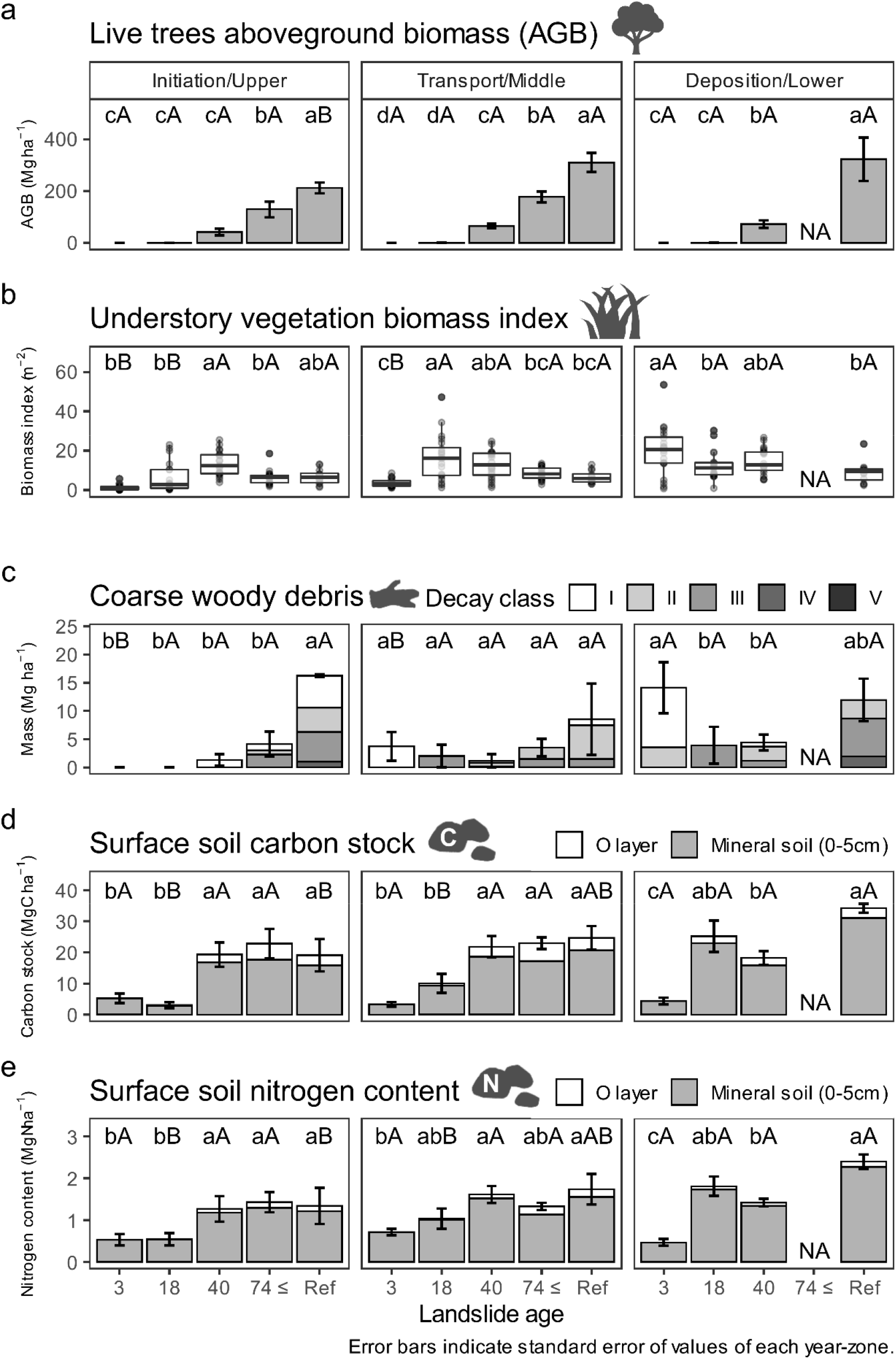
Chronological changes in each forest ecosystem component in each slope zone. (a) The chronological changes in live tree AGB in each slope zone. (b) The chronological changes in the biomass indices of understory vegetation in each slope zone. (c) The chronological changes in coarse woody debris (CWD) mass in each slope zone. (d) The chronological changes in surface soil carbon stock in each slope zone. White and gray bars indicate the O layer and mineral soil carbon stock, respectively. (e) The chronological changes in surface soil nitrogen content in each slope zone. White and gray bars indicate the O layer and mineral soil nitrogen content, respectively. Error bars in a, c, d, and e indicate standard errors of each forest ecosystem component of each combination of landslide age and zone. Different lowercase letters above the bars indicate significant differences between landslide ages within each zone (p < 0.05) according to multiple comparisons. Different capital letters above the bars indicate significant differences between zones within each landslide age (p < 0.05) according to multiple comparisons. The data for landslide age 3 are the same data shown in Table 2.

### Understory vegetation

Each zone had unique patterns of chronological changes in the biomass indices of understory vegetation. Unimodal distributions with peaks at 18 and 40 years after landslides were observed in the transport and initiation zones, respectively (Figure 2b, left and center). Biomass indices in the deposition zone were largest at 3 years after landslides and monotonically decreased with years elapsed (Figure 2b, right). In addition, the biomass indices 3 years after landslides in the deposition zone were significantly larger than those in the initiation and transport zones. The biomass indices 18 years after landslides in the transport zone were significantly larger than those in the initiation zone.

### CWD

The chronological changes in CWD mass also had unique patterns dependent on zone. The CWD mass at the initiation zone was largest in the reference stands and monotonically increased with landslide age (Figure 2c, left). U-shaped chronological changes in CWD mass with peaks at 3 years after landslides and reference stands were observed in the deposition zone (Figure 2c, right). A similar pattern was also observed in the transport zone, although there were no significant differences with time after landslide (Figure 2c, center). In addition, the CWD mass 3 years after landslides was significantly larger in the deposition zone than in the initiation and transport zones. Chronological changes in CWD decay class compositions were also different among zones (Figure 2c). CWD in decay classes III, IV, or V was observed in the transport and deposition zones in areas 18 or more years after landslides; however, these classes were not observed in the initiation zone unless it was more than 74 years after landslides.

### Surface soils

The recovery rates of surface soil CN were different among the zones. In the initiation and transport zones, surface soil CN 40 and over 74 years after landslides were not significantly different from those in the reference stands; thus, surface soil CN recovered to their reference levels within 40 years after landslides (Figure 2d and e, left and center). In the deposition zone, the surface soil CN 18 years after landslides were not significantly different from those in the reference stands, thus recovering to their reference levels within 18 years after landslides (Figure 2d and e, right). Additionally, the surface soil CN 18 years after landslides were significantly larger in the deposition zone than in the initiation zone. Both carbon stock and nitrogen content showed similar patterns of temporal changes.

### Live tree species composition

The species composition of live trees was significantly different among landslide ages and showed different trends among zones. Regardless of zone, live tree species composition was significantly different among the landslide ages according to PERMANOVA (Figure 3c). These differences were also visually observed in the NMDS ordination result (Figure 3a). Plots at landslide age 40 were characterized by shade-intolerant species such as *Betula platyphylla* and *Alnus hirsuta*; however, plots at landslide age ≥ 74 and the reference were characterized by intermediate and shade tolerant species such as *Acer pictum, Quercus crispula*, and *Tilia japonica* (Figure 3a and b). These tendencies were also recognized the dominance rate of the basal area (Figure 3c). Desiccation-intolerant species tended to be most dominant in the order of the deposition, transport, and initiation zones (Figure 3a, b, and d). Regardless of landslide age, the NMDS results showed that plots in the deposition zone were characterized by desiccation-intolerant species such as *Alnus hirsuta, Carpinus cordata, Ulmus davidiana* var. *japonica*, and *Cercidiphyllum japonicum* (Figure 3a and b). Desiccation-intolerant species accounted for more than 50% of the total basal area in the deposition/lower zone regardless of landslide age (Figure 3d). The slope angle, soil moisture content, and surface soil nitrogen content were significantly correlated to the ordination results (Figure 3a; Appendix 4). The surface soil nitrogen content and O layer depth were suggested to mainly indicate the environmental differences among landslide ages, while the soil moisture content and slope angle were suggested to mainly indicate the environmental differences among zones.

**Figure 3.**
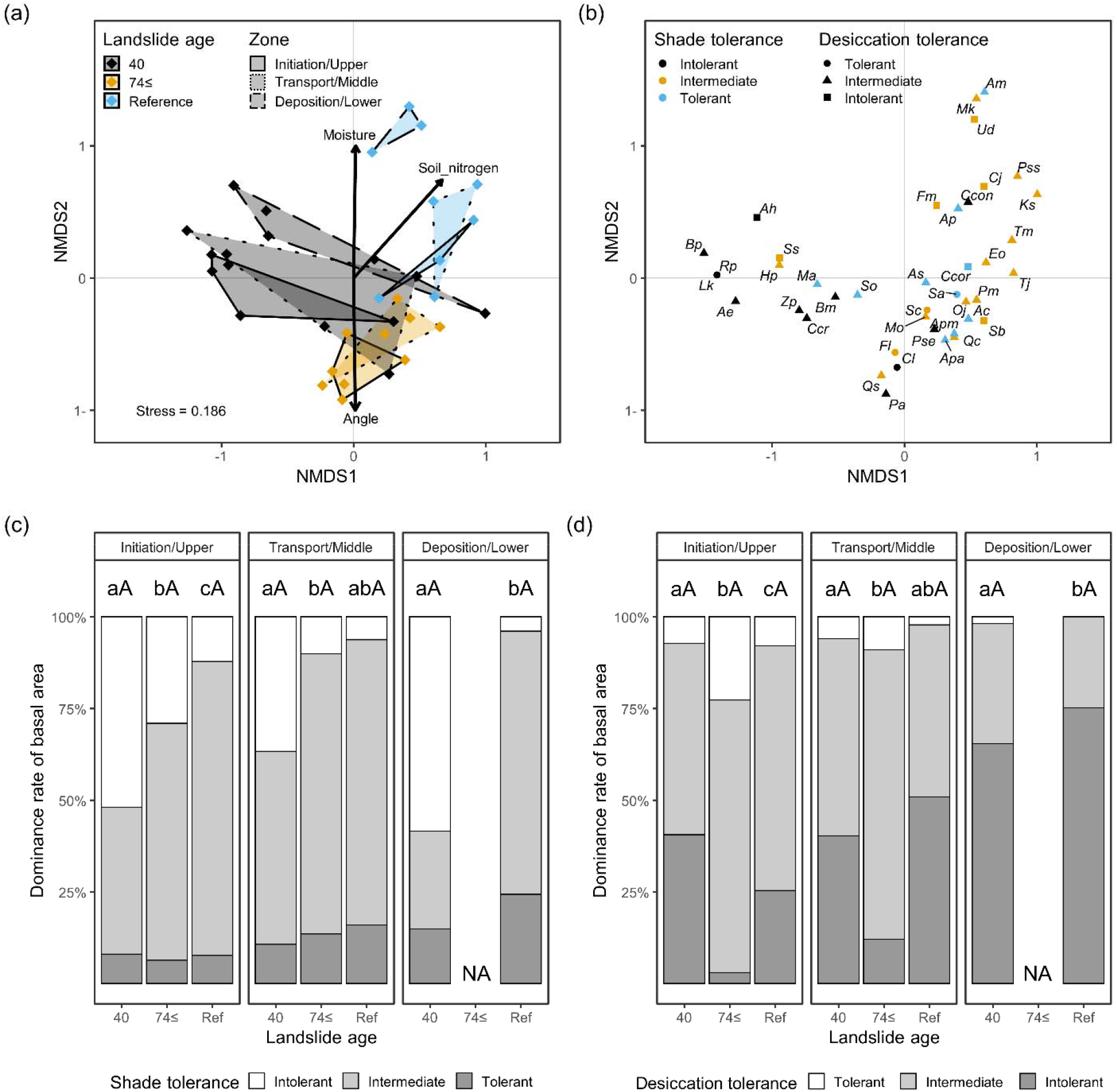
Live tree species composition based on basal area. Ordination analysis (NMDS) of live trees for (a) plots and environmental variables (only significant variables (p < 0.05) shown) and for (b) species. Colors in (a) indicate landslide ages, and line types in (a) indicate zones. Colors in (b) indicate species shade tolerance, and point shapes in (b) indicate species desiccation tolerance. For full species names, please see Appendix 3. (c) Dominance rate of basal area classified by shade tolerance. (d) Dominance rate of the basal area classified by desiccation tolerance. Different lowercase letters above the bars indicate significant differences in species composition between landslide ages within each zone (p < 0.05) according to PERMANOVA and multiple comparisons. Different capital letters above the bars indicate significant differences in species composition between zones within each landslide age (p < 0.05) according to PERMANOVA and multiple comparisons.

### Understory vegetation species composition

Understory vegetation species composition was significantly different among the landslide ages and zones according to PERMANOVA and NMDS (Figure 4a, b, c, and d). Quadrats with younger landslide ages were characterized by open land species and species that favor dry environments; however, quadrats with older landslide ages were characterized by forest species (Figure 4a, b, c, and d). The dominance rate of vegetation cover also showed a decrease in open land species and an increase in generalist and forest species with increasing landslide age (Figure 4e). In particular, the dominance rate of forest species cover increased more rapidly, and the dominance rate of open land species cover decreased more rapidly in the deposition zone than in the initiation zone (Figure 4e). The dominance rate of forest species cover in the deposition zone 18 years after landslides was 43%, which was approximately three times that in the initiation zone, i.e., 14% (Figure 4e). The dominance rate of open land species cover in the deposition zone 18 after landslides was 16%; however, that in the initiation zone was 26% (Figure 4e). Regardless of landslide ages (except for age 3), quadrats in the deposition/lower zones were characterized by species that favor moist environments (Figure 4c and d), and the dominance rate of this type of species cover was larger in the deposition/lower zone than in the initiation/upper and transport/middle zones (Figure 4f). AGB, slope angle, soil moisture content, O layer depth, and surface soil nitrogen content were significantly correlated with the ordination results (Figure 4a, b, c; Appendix 5). AGB, surface soil nitrogen content, and O layer depth mainly indicated the environmental differences among landslide ages. Additionally, the soil moisture content and slope angle mainly indicated the environmental differences among zones.

**Figure 4.**
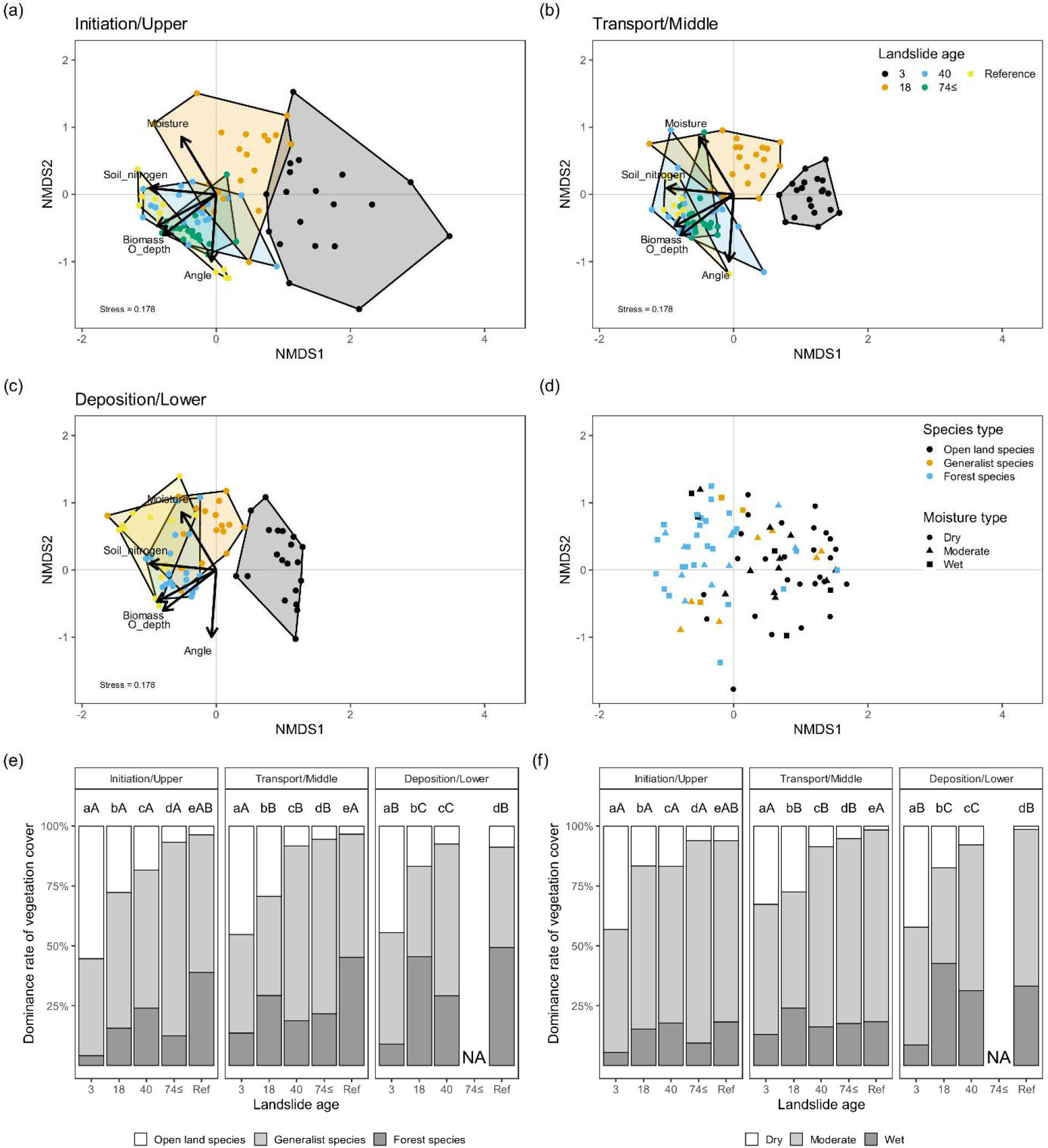
Understory vegetation species composition based on vegetation cover. NMDS results of understory vegetation for (a) quadrats in the initiation/upper zone, (b) quadrats in the transport/middle zone, (c) quadrats in the deposition/lower zone, and (d) species. Arrows in (a), (b), and (c) indicate the directions of the greatest environmental gradient of each environmental variable (only significant variables (p < 0.05) shown). Notably, the same ordination result is divided and displayed by zone in (a), (b), and (c). The colors in (a), (b), and (c) indicate landslide age. Colors in (d) indicate species type, and point shapes in (d) indicate species moisture type. (e) Dominance rate of herbaceous species vegetation cover classified by species type. (f) Dominance rate of herbaceous species vegetation cover classified by moisture type. Different lowercase letters above the bars indicate significant differences in species composition between landslide ages within each zone (p < 0.05) according to PERMANOVA and multiple comparisons. Different capital letters above the bars indicate significant differences in species composition between zones within each landslide age (p < 0.05) according to PERMANOVA and multiple comparisons.

## DISCUSSION

We analyzed 74 years of changes in four forest ecosystem components (i.e., live trees, understory vegetation, CWD, and surface soils) in three zones generated by landslides. Our results illustrate that zone-dependent patterns of chronological changes in each forest ecosystem component, i.e., the succession and recovery of forest ecosystems, occurred most rapidly in the deposition zone, followed by that in the transport zone and then the initiation zone. As we hypothesized, the accumulation of live tree AGB was slower in the initiation zone than in the other zones, as disturbance legacies in the initiation zone were minimal and surface soils were unstabilized due to the higher slope angle. The chronological changes in the amounts of understory vegetation were determined by the difference in the amounts of disturbance legacies just after a landslide and the timing of subsequent canopy closure, which was dependent on zones. For the same reason, the succession of understory vegetation species composition occurred most rapidly in the deposition zone, followed by in the transport zone and then the initiation zone. The chronological changes in the amounts of CWD were determined by the difference in the amounts of disturbance legacies just after a shallow landslide and mortality through stand developments. Furthermore, zone-dependent chronological changes in live trees, understory vegetation, and CWD influenced subsequent litter production and determined the recovery rates of surface soil CN. In contrast to our hypothesis, the succession of live tree composition was not significantly different among the zones. Low slope angles and disturbance legacy richness facilitated AGB accumulation; however, they did not affect the successional rate of species composition.

### Live trees

Live tree AGB recovery was facilitated by the low slope angle and disturbance legacy richness. The slope angle was greatest in the initiation zone, followed by the transport zone and then the deposition zone (Table 1). More surface soils are eroded and lost on steep slopes due to rainfall (Warrington and others 1989; Martínez-Murillo and others 2013). Thus, surface soils in the initiation zone were less stable than those in the transport and deposition zones, and it was more difficult for tree seedlings to establish, particularly during several years just after landslides (Buma and Pawlik 2021; Guariguata 1990). Moreover, needle ice forms and melts in soils during the winter season in the cool-temperate climate regions where our study sites were located, and this process destabilizes surface soils and facilitates soil erosion (Prosser and others 2000). Thus, this factor may have been a why live tree recovery in the present study was slower than that in warmer climate regions as reported in previous studies (Schomakers and others 2017; Joshi and Garkoti 2021). Disturbance legacies (i.e., undisturbed vegetation patches and CWD) were richest in the deposition zone, followed by in the transport zone and then the initiation zone (Table 2) because they were transported from the upper part of slopes when shallow landslides occurred (Restrepo and others 2009; Walker and others 2009). The amounts of CWD and understory vegetation 3 years after landslides were much larger in the deposition zone than in the initiation and transport zones (Table 2), which directly indicated that the amounts of disturbance legacy were larger in the deposition zone, although the understory vegetation in this zone 3 years after landslides contained some newly dispersed and established vegetation after the landslides. At disturbance legacy-rich sites, live trees can regenerate by advanced seedlings on undisturbed vegetation patches, buried seeds, or sprouts (Guariguata 1990). Freund and others (2021) also suggested that canopy heights are higher for 25 years after landslides at sites that include residual vegetation. Therefore, the accumulation rates of live tree AGB were highest (Figure 2a) in the deposition zone, followed by the transport zone and then the initiation zone. Supplementary, live tree recovery after shallow landslides was relatively slower than that of the other disturbance types. Please see Appendix 7 for a detailed discussion.

### Understory vegetation

Disturbance legacy richness, soil angles, and timing of canopy closure associated with stand development determined the chronological patterns of understory vegetation amounts dependent on zones. The understory vegetation biomass index 3 years after landslides was larger in the deposition zone than in the other two zones (Table 2; Figure 2b, landslide age 3), which could be attributed to the richness of undisturbed vegetation patches, buried seeds, rhizomes and the lower slope angle, as discussed above. In particular, species that reproduce via rhizomes, such as *Sasa nipponica* (Makino) Makino et Shibata and *Petasites japonicus* subsp. *Giganteus*, may have assisted in rapid vegetation recovery (Makita and others 1993; Hind and Kay 2006). However, in the initiation and transport zones, seed dispersal and establishment were needed for vegetation recovery, and surface soils were exposed to soil erosion, as mentioned above; thus, the vegetation recovery was slower than that in the deposition zone (Buma and Pawlik 2021). The canopy closure associated with the increase in live tree AGB caused the understory vegetation biomass indices to finally decrease regardless of zone (Figure 2b). Generally, dense canopy cover results in sparse or no understory vegetation (Sun and others 2016); thus, the amount of understory vegetation gradually decreases with stand development (Jules and others 2008). Live tree AGB accumulated more slowly in the initiation zone than in the other two zones (Figure 2a), which suggests that canopy closure occurred later in the initiation zone. Therefore, biomass indices began to decline 40 to 74 years after landslides in the initiation zone, which was later than that in the deposition zone.

### CWD

The chronological patterns of the CWD amounts were determined mainly by the loss and accumulation of CWD due to landslides and the additional input of CWD through subsequent stand development. Once landslides occurred, live trees and CWD on the slope were transported to and accumulated at the lower part of the slope (landslide mortality, which partially includes carryover). In fact, our data also showed an uneven distribution of CWD on the slopes just after landslides, i.e., more CWD in the deposition zone and less CWD in the initiation and transport zones (Table 2; Figure 2c, landslide age 3). These unevenly distributed CWD was decomposed, which likely resulted in differences in the amounts of decayed CWD (decay class III, IV, or V) among the zones 18 and 40 years after landslides (Figure 2c). Regardless of zone, the amount of CWD eventually tended to increase due to the increase in mortality by self-thinning through stand development (Yoda and others 1963). Therefore, the combination of mortality due to landslides and the CWD input from the new stand determined the chronological patterns of the CWD amount in each zone. Thus, the chronological changes in CWD in the initiation zone showed only CWD input from the new stand, while those in the transport zone showed a few CWD input by landslides and CWD input from the new stand; finally, those in the deposition zone showed a lot of CWD input by landslides and CWD input from the new stand, which was a “U-shaped temporal pattern” (Spies and others 1988). U-shaped temporal patterns were also reported in previous studies focused on other disturbance types, such as wildfire, windthrow, and harvest (Brassard and Chen 2008; Krankina and Harmon 1995; Spies and others 1988). Unlike other disturbance types, landslides generate chronological patterns of CWD amounts that can be completely different within disturbed areas.

### Surface soil CN

The recovery rates of surface soil CN dependent on zones were determined by the difference in slope angles and the amounts of litter, which was induced by both the amounts of CWD just after the landslides and the recovery rates of understory vegetation and live trees. As we have discussed above, more soils eroded on the upper part of the slopes; thus, soil development was relatively slow in the upper part of the slopes. Additionally, the accumulation rates of understory vegetation and live tree AGB were fastest in the deposition zone, followed by the transport zone and then the initiation zone (Figure 2a, b, c). These results suggest that the amounts of litter of both understory vegetation and live trees were also greatest in the deposition zone, followed by the transport zone and then the initiation zone. In forest ecosystems, litter fall is one of the main sources of soil carbon and nitrogen (Duan and others 2020). Additionally, the amounts of CWD just after the shallow landslide events were also greatest in the deposition zone, followed by the transport zone and then the initiation zone. This leaf and wood litter is decomposed and transferred to surface soils. Thus, the surface soil CN recovered to the level of the reference stands faster in the deposition zone (18 years), followed by the transport zone (18-40 years) and then the initiation zone (40 years). Wicke and others (2003) also reported that the accumulation of soil nutrients occurred more rapidly at the lower part of landslides within 20 years, although they did not evaluate the recovery times to the level of reference stands.

Regardless of zone, surface soil CN recovered to the level of reference stands faster than the recovery of live tree AGB, which took more than 74 years. Previous studies have also reported a faster recovery of surface soil CN than of live trees (Pandey and Singh 1985; Reddy and Singh 1993). Because soil nitrogen is one of the most important factors for plant succession and tree growth (Shiels and others 2008; Walker and others 2009), surface soil CN recovery rates might influence deeper soil CN recovery and long-term (over 100 years) live tree AGB recovery, which was beyond the scope and timescale of the present study.

### Live tree and understory vegetation species composition

Landslide age- and zone-dependent environmental conditions are suggested to affect live tree and understory vegetation species composition. Higher surface soil nitrogen content and O layer depth (Figure 3a and 4a, b, and c), which were greatly correlated with landslide ages and partly correlated with zones (Figure 2e; Table 1), resulted in shade-tolerant species and forest species dominance among the live trees and understory vegetation, respectively. In general, soil nitrogen content and O layer depth (significant only in the understory) positively correlate with soil development and fertility (Pritchett and Fisher 1987). Shade-tolerant species are generally large-seeded species (Reich and others 1998). Relatedly, Jager and others (2015) suggested that large-seeded shade-tolerant species favor high-fertility soils. Thus, the soil nitrogen content and O layer depth coupled with years elapsed after shallow landslides would facilitate shade-tolerant species dominance. Increasing AGB positively correlated canopy closure with landslide age, which strongly affected the understory vegetation species composition. Live tree AGB recovery occurred most rapidly in the deposition zone, followed by the transport zone and then the initiation zone, due to the low soil erosion levels and rich biological legacies in the deposition zone, as discussed above. Thus, species composition succession of understory vegetation occurred fastest in the deposition zone, followed by the transport zone and then the initiation zone. Significant effects of soil moisture content and slope angle (Figure 3a; Figure 4a, b, c, and d), which were higher and smaller in order of the deposition, transport, and initiation zones (Table 1), suggested the presence of zone effects on the species composition of live trees and understory vegetation, although there were no significant differences in live tree species composition among the zones. In general, the deposition zones were located along or above streams (Appendix 1) because debris generated by landslides continues downward until it reaches flat terrain or a valley. Thus, forests such as riparian forests then developed, and species that favor moist environments were dominant in the deposition zones.

## DATA AVAILABILITY

Data and code are available on Dryad: https://datadryad.org/stash/share/0cVLSf5uOq-CE3xeuDUIy7FbFgUrMibs85Ub57uR-Y8

## ACKNOWLEDGMENTS

We thank the members of the Ecosystem Management Laboratory for helping with the field survey and for their helpful discussions about our study. We thank Ms. Yuriko Horiuchi, Ms. Akari Kimura, Dr. Takahiro Inoue, Dr. Hayato Maruyama and Dr. Toshihiro Watanabe for helping with the soil analysis. We thank Dr. Hajime Sato for allowing us to use data of historical landslides, Dr. Satoshi N. Suzuki and Dr. Kazuhiro Kawamura for helpful advice about statistical analysis, and Dr. Takashi Yamada for valuable advice on the study design. We would like to thank Atsuma town, Hidaka town, Biratori town, and Hekiun Ranch Co., Ltd., for allowing us to study these sites.

