## Appendix for "Long-term recoveries of forest ecosystem components after shallow landslides were asynchronized among slope positions"

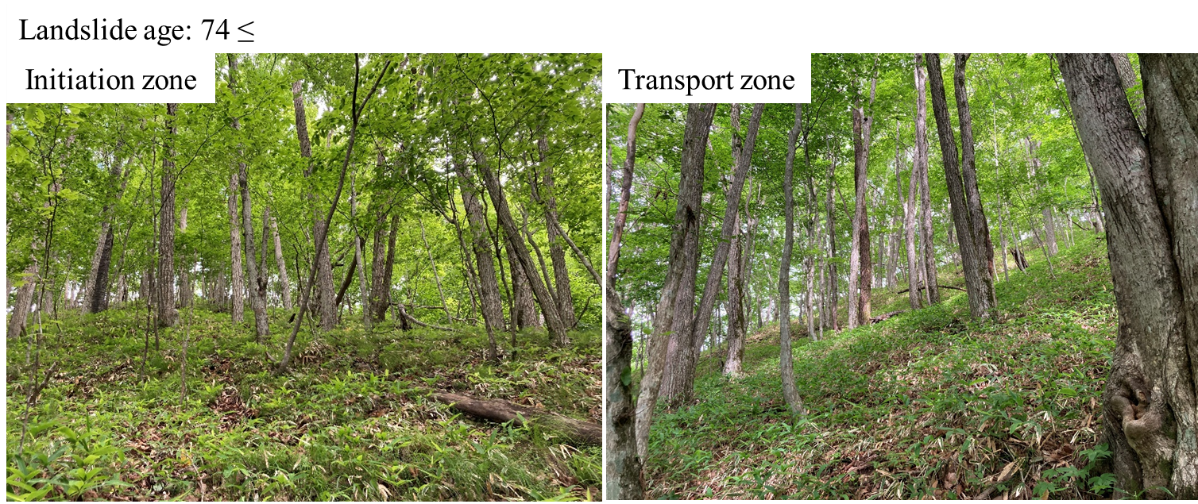
**Appendix 1. Photographs of each landslide age and zone**

**Figure S1.1. Photographs of forests with landslide ages over 74.**

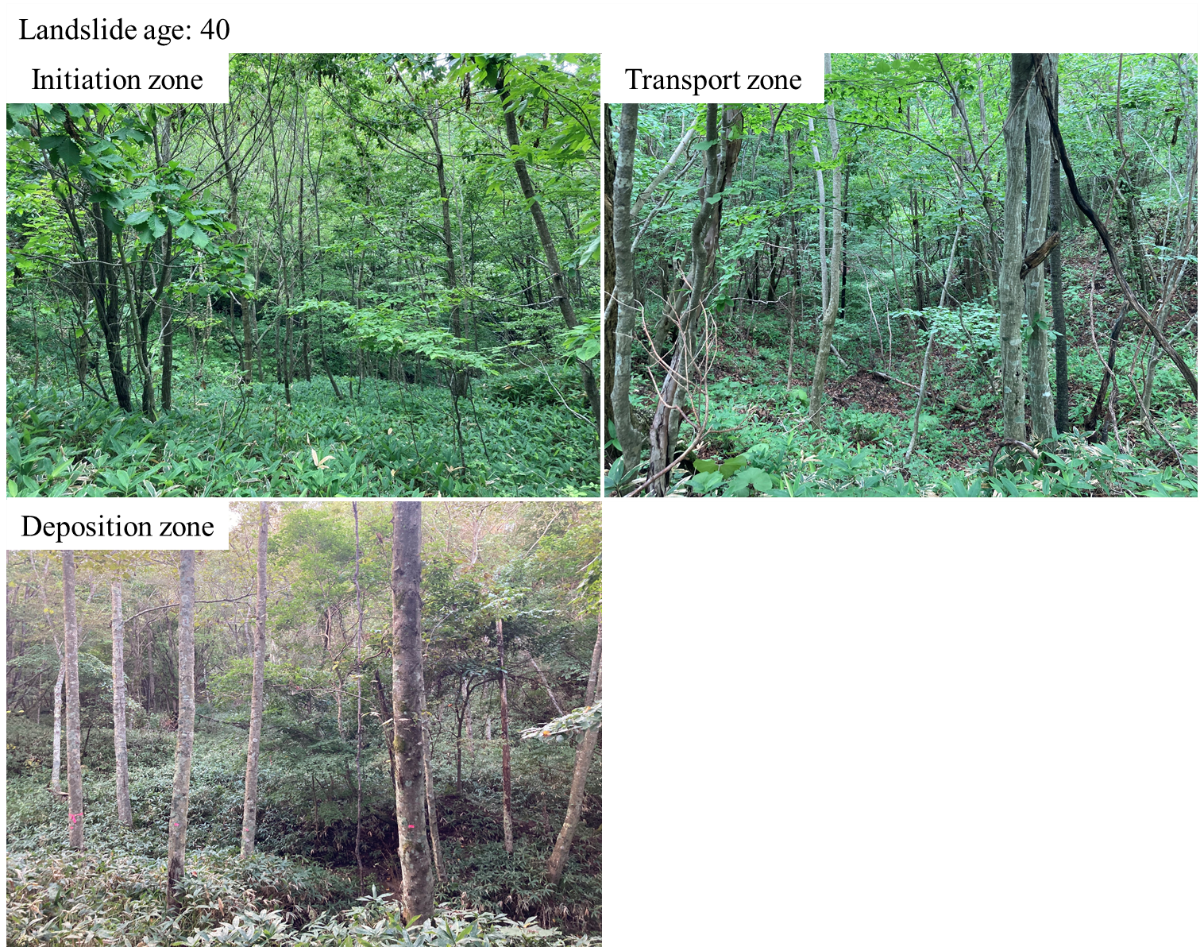

**Figure S1.2. Photographs of forests with a landslide age of 40.**

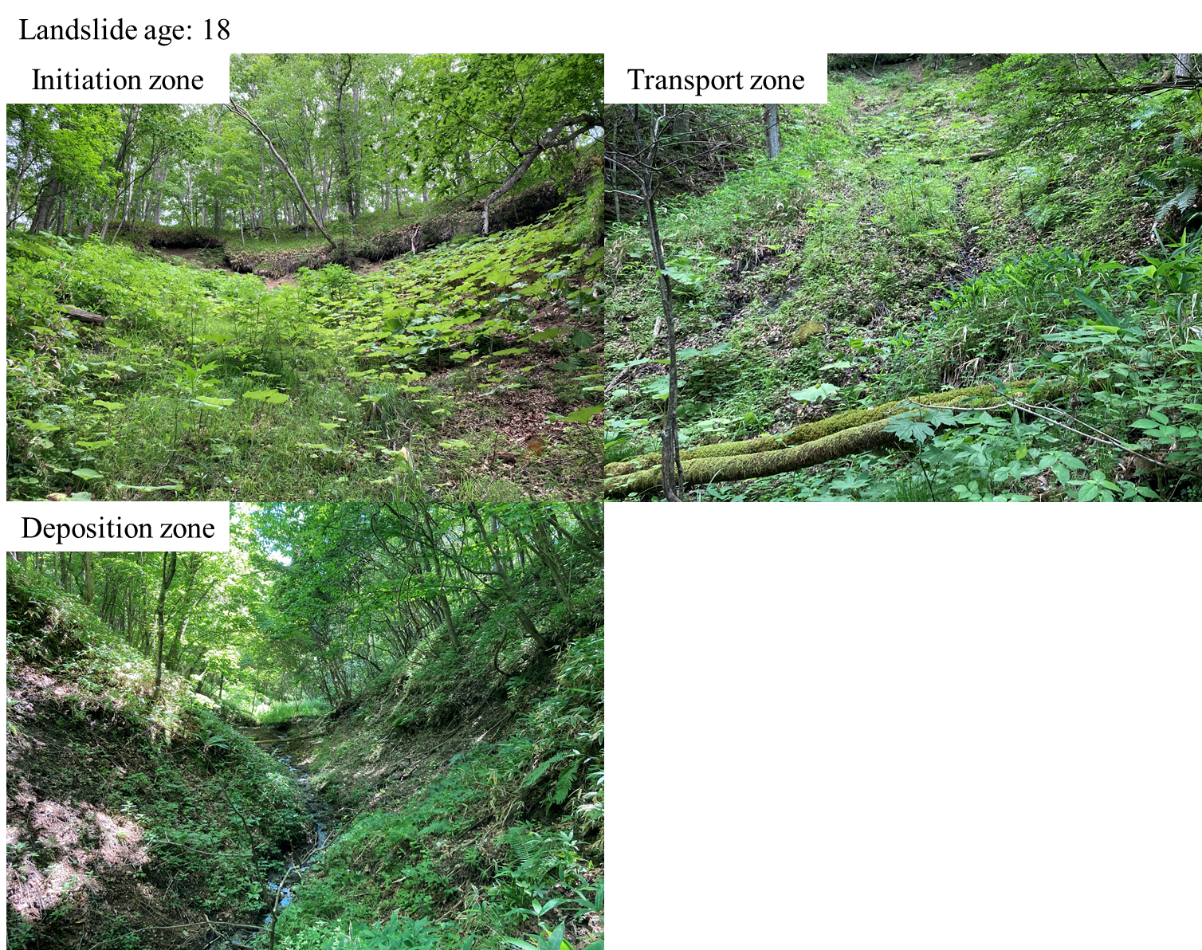
**Figure S1.3. Photographs of forests with a landslide age of 18.**

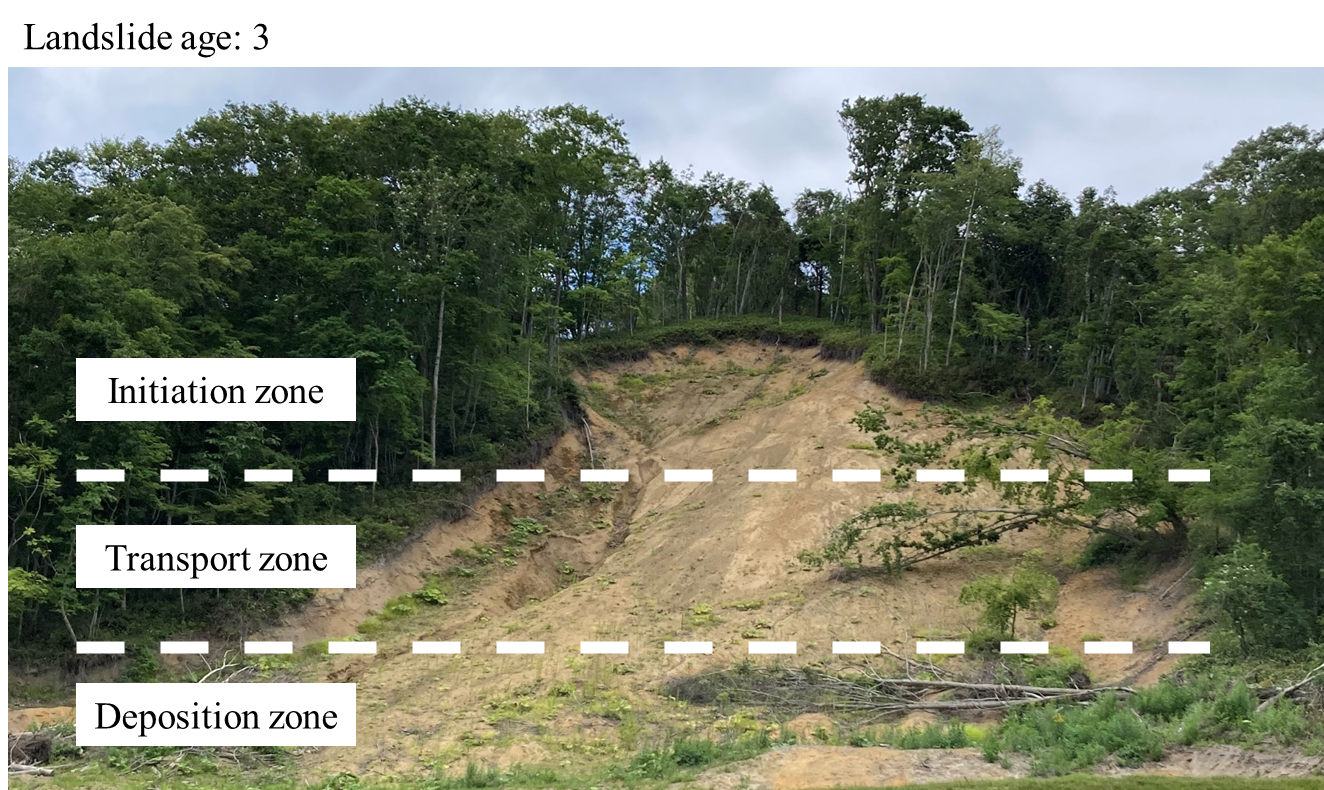

**Figure S1.4. Photographs of forests with a landslide age of 3.**

**Appendix 2. Calculations of live tree aboveground biomass and CWD mass**

Live tree aboveground biomass (*AGB*) was calculated as *AGB* = *V* × *BEF* × *D*, where *V* is the volume of the stem, *BEF* is the species-specific biomass extension factor (GIO 2016), and *D* is the species-specific volume density (GIO 2016). The stem volumes of conifers were estimated by using regional derived allometric equations for each species (Japanese Forestry Agency planning division 1970). Broadleaved tree species were classified into three groups, and the stem volumes of the broadleaved trees were estimated by using allometric equations for each group (Maezawa and others 1968).

Downed log mass (*M_DownedLogs_*) was calculated as *M_DownedLogs_* = *L*/12 × {5*A_b_* + 5*A_u_* + 2√*A_b_A_u_*} × *D*, where *L* is the length of the downed log, *A_b_* is the cross-sectional area of the larger end of the downed log, *A_u_* is the cross-sectional area of the smaller end of the downed log, and *D* is the specific value of the volume density for each decay class and tree species (Ugawa and others 2012). The method of the approximation to a conic-paraboloid (Fraver and others 2007) was applied for the calculation of the downed log volume. No downed logs with roots were observed, so the mass of dead roots was not calculated.

Snag mass (*M_Snags_*) was calculated as follows: for defective snags, first, we estimated the original height of snags as 1/*L_1_* = 1/(1.21 × *DBH*^0.946^) + 1/19.5, where *L_1_* is the height of the snag when the full body remained and *DBH* is the diameter at breast height (Goto and others 2003). The volume of the snag when the full body remained (*V_1_*) was calculated by *DBH* and *L_1_* by using allometric equations for live tree stem volume (Japanese Forestry Agency planning division 1970; Maezawa and others 1968). The snag volume (*V_Snags_*) was calculated as *V_Snags_* = [*V_1_* – *V_1_* × {(*L* – *L_1_*)/*L*}^3^]. For intact snags, the snag volume (*V_Snags_*) was calculated by the observed height and DBH of the snag by using allometric equations. Last, snag mass (*M_Snags_*) was calculated as *M_Snags_* = *V_Snags_* × (1 + *R*) × *D*, where *R* is the species-specific ratio of the root to the stem (GIO 2016) and *D* is the volume density of the value of the volume density for each decay class and tree species (Ugawa and others 2012).

**Appendix 3. Shade tolerance, moisture type, and abbreviations of the live tree species**

**Table S3. Shade tolerance, moisture type, and abbreviations of the live tree species**

| Scientific name | Abbreviation | Shade tolerance | Desiccation tolerance |
| --- | --- | --- | --- |
| *Abies sachalinensis* | *As* | Tolerant | Intermediate |
| *Acer cissifolium* | *Ac* | Tolerant | Intermediate |
| *Acer japonicum* | *-* | Tolerant | Intermediate |
| *Acer miyabei* | *Am* | Tolerant | Intermediate |
| *Acer palmatum* var. *amoenum* | *Apa* | Tolerant | Intermediate |
| *Acer palmatum* var.*mastumurae* | *Apm* | Tolerant | Intermediate |
| *Acer pictum* | *Ap* | Tolerant | Intermediate |
| *Actinidia arguta* | *-* | Intermediate | Intermediate |
| *Alnus hirsuta* | *Ah* | Intolerant | Intolerant |
| *Aralia elata* | *Ae* | Intolerant | Intermediate |
| *Betula maximowicziana* | *Bm* | Intolerant | Intermediate |
| *Betula platyphylla* | *Bp* | Intolerant | Intermediate |
| *Carpinus cordata* | *Ccor* | Tolerant | Intolerant |
| *Carpinus laxiflora* | *Cl* | Intolerant | Tolerant |
| *Castanea crenata* | *Ccr* | Intolerant | Intermediate |
| *Cercidiphyllum japonicum* | *Cj* | Intermediate | Intolerant |
| *Cornus controversa* | *Ccon* | Intolerant | Intermediate |
| *Euonymus oxyphyllus* | *Eo* | Intermediate | Intermediate |
| *Fraxinus lanuginosa* | *Fl* | Intermediate | Tolerant |
| *Fraxinus mandshurica* | *Fm* | Intermediate | Intolerant |
| *Hydrangea paniculata* | *Hp* | Intermediate | Intermediate |
| *Kalopanax septemlobus* | *Ks* | Intermediate | Intermediate |
| *Larix kaempferi* | *Lk* | Intolerant | Tolerant |
| *Lespedeza bicolor* | *-* | Intermediate | Intermediate |
| *Magnolia kobus* var. *borealis* | *Mk* | Intermediate | Intermediate |
| *Magnolia obovata* | *Mo* | Intermediate | Intermediate |
| *Morus australis* | *Ma* | Tolerant | Intermediate |
| *Ostrya japonica* | *Oj* | Intermediate | Intermediate |
| *Phellodendron amurense* | *Pa* | Intolerant | Intermediate |
| *Prunus maximowiczii* | *Pm* | Intermediate | Intermediate |
| *Prunus sergentii* | *Pse* | Intolerant | Intermediate |
| *Prunus ssiori* | *Pss* | Intermediate | Intermediate |
| *Quercus crispula* | *Qc* | Intermediate | Intermediate |
| *Quercus serrata* | *Qs* | Intermediate | Intermediate |
| *Robinia pseudoacacia* | *Rp* | Intolerant | Tolerant |
| *Salix bakko* | *Sb* | Intermediate | Intolerant |
| *Salix integra* | *-* | Intolerant | Intermediate |
| *Salix sachalinensis* | *Ss* | Intermediate | Intolerant |
| *Sorbus alnifolia* | *Sa* | Tolerant | Tolerant |
| *Sorbus commixta* | *Sc* | Intermediate | Tolerant |
| *Staphylea bumalda* | *-* | Intermediate | Intermediate |
| *Styrax obassia* | *So* | Tolerant | Intermediate |
| *Tilia japonica* | *Tj* | Intermediate | Intermediate |
| *Tilia maximowicziana* | *Tm* | Intermediate | Intermediate |
| *Toxicodendron trichocarpum* | *-* | Intermediate | Intermediate |
| *Ulmus davidiana* var. *japonica* | *Ud* | Intermediate | Intolerant |
| *Viburnum dilatatum* | *-* | Intermediate | Intermediate |
| *Viburnum furcatum* | *-* | Intermediate | Intermediate |
| *Zanthoxylum piperitum* | *Zp* | Intolerant | Intermediate |

Species whose abbreviation is “-” were not included in the NMDS ordination.

**Appendix 4. Environmental gradient contours** **of the NMDS of the live tree species composition**

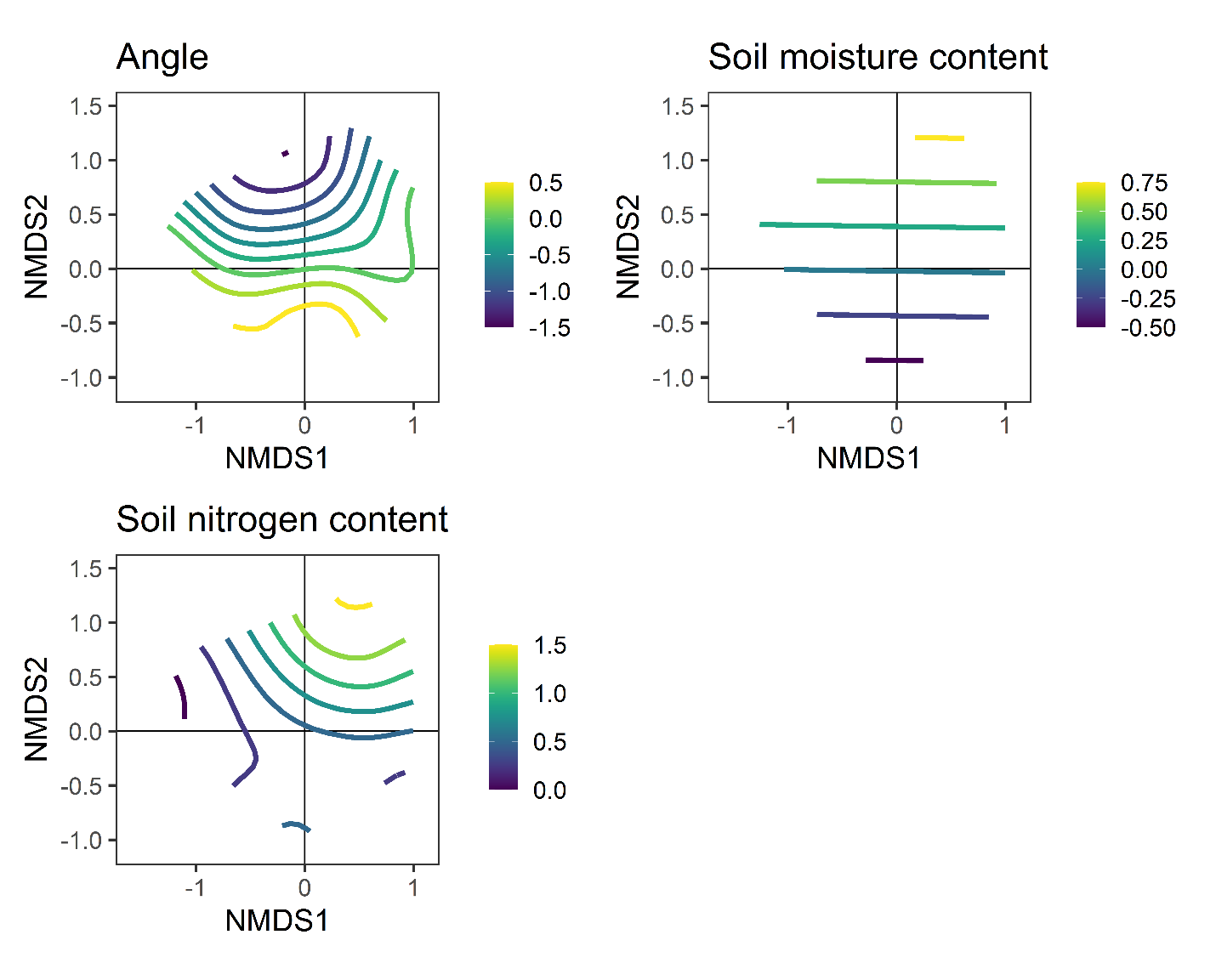

**Figure S4. Contour lines of the environmental variables of the NMDS of live tree species composition are shown in Figure 3a.**

**Appendix 5. Environmental gradient contours of the NMDS of the understory vegetation species composition**

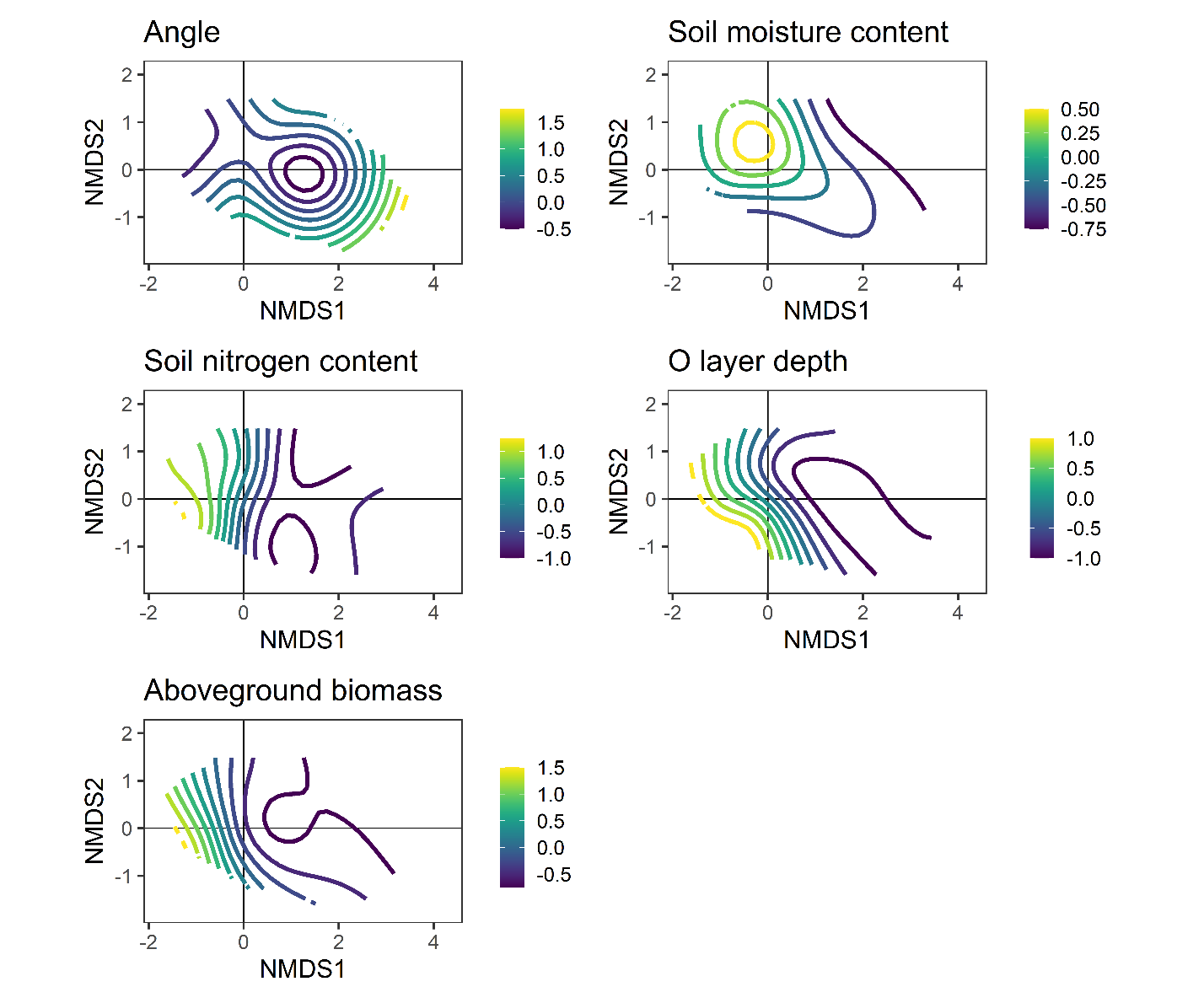

**Figure S5. Contour lines of the environmental variables of the NMDS of the understory vegetation are shown in Figure 4a, b, and c.**

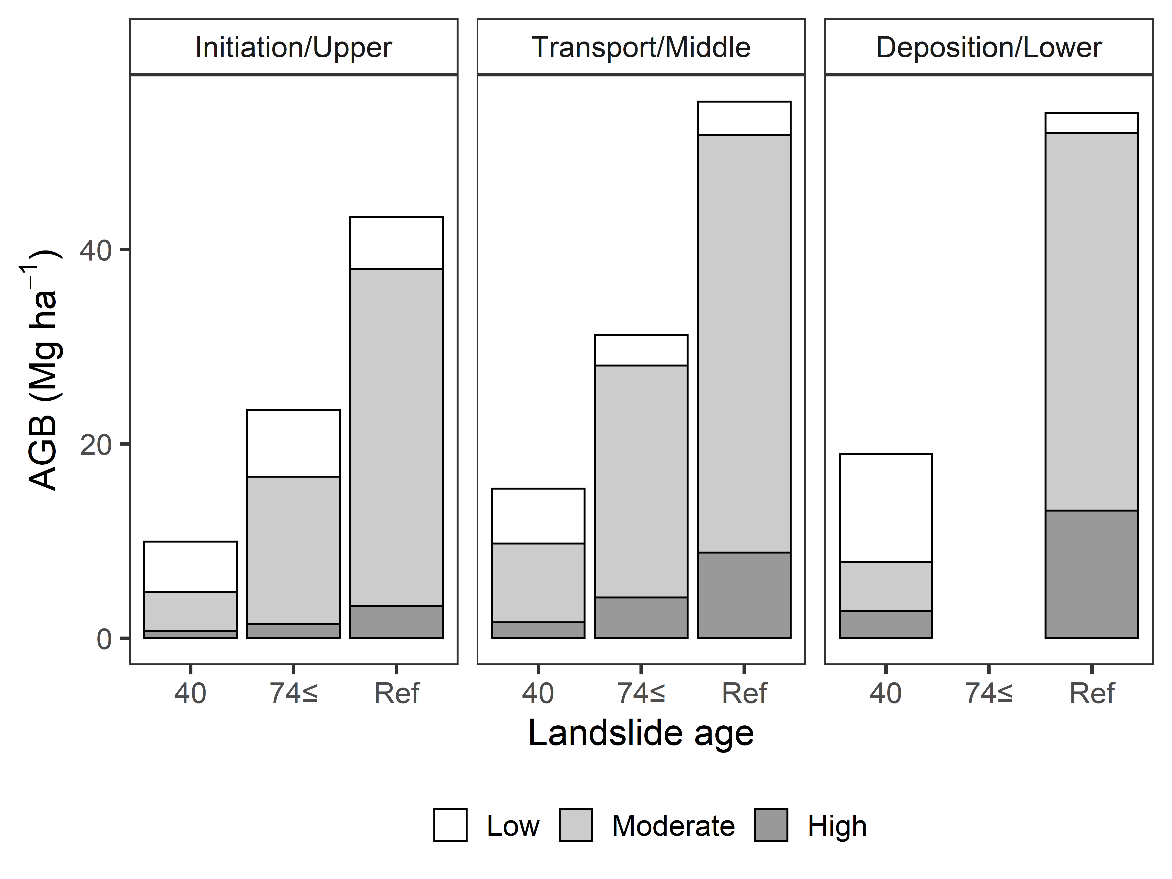
**Appendix 6. Aboveground biomass by shade tolerance and moisture type**

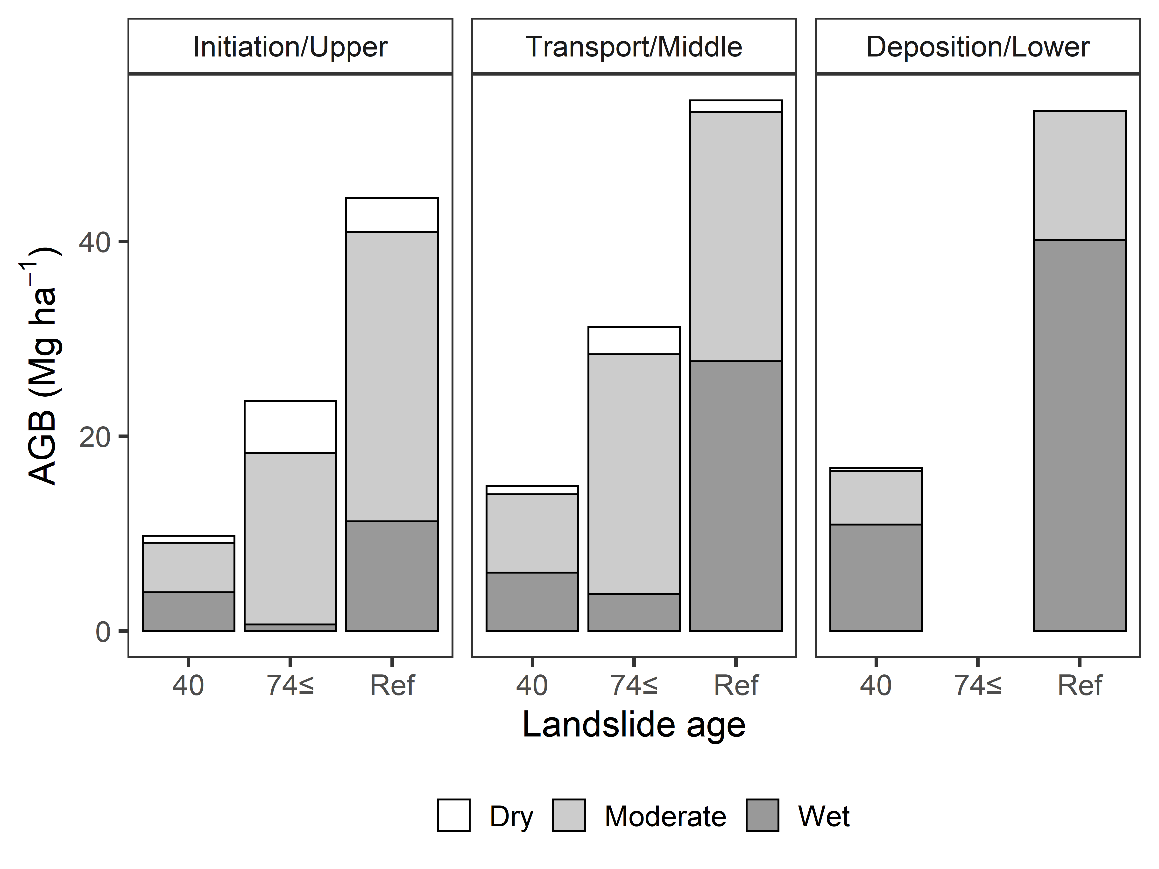
**Figure S6.1. Aboveground biomass by shade tolerance for each landslide age and zone.**

**Figure S6.2. Aboveground biomass by moisture type for each landslide age and zone.**

**Appendix 7. The mass and carbon and nitrogen concentrations of the O layer and mineral soil samples**

The O layer was divided into L, FH, and twigs in Figure S7.1.
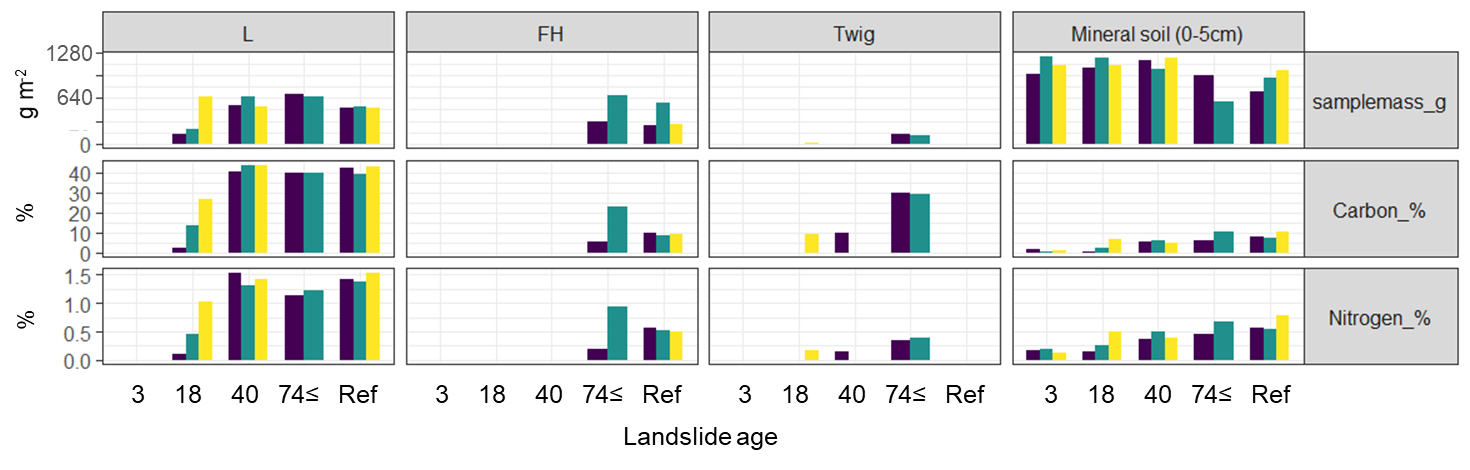

**Figure S7.1. The mass and carbon and nitrogen concentrations of the O layer and mineral soil samples**

**Appendix 8. Supplementary discussion: Comparisons of recovery rates among different disturbance types.**

Live tree recovery in the initiation and transport zones (i.e., disturbance legacy-poor zones) resembled primary succession; thus, the recovery rates were also lower than those after the other disturbances. In the present study, live tree AGB in the legacy-poor zones recovered 57-59% of those in the reference stands within 74 years. In contrast, Vijayakumar and others (2016) reported that the AGB of live trees recovered to 84-100% of those in reference stands within 61-90 years after wildfires. Hotta and others (2020) and Suzuki and others (2019) reported that the AGB of live trees recovered to 82% and 71% of those in reference stands within 64 and approximately 55 years after windthrow, respectively. In the case of wildfires and windthrow, some advanced seedlings survived, and CWD was left intact after disturbance events; thus, live trees recovered faster than after the landslide.
